# A redox-shifted fibroblast subpopulation emerges in the fibrotic lung

**DOI:** 10.1101/2023.09.23.559128

**Authors:** Patrick A. Link, Jeffrey A. Meridew, Nunzia Caporarello, Ashley Y. Gao, Victor Peters, Gordon B. Smith, Mauricio Rojas, Daniel J. Tschumperlin

## Abstract

Idiopathic pulmonary fibrosis (IPF) is an aggressive and thus far incurable disease, characterized by aberrant fibroblast-mediated extracellular matrix deposition. Our understanding of the disease etiology is incomplete; however, there is consensus that a reduction-oxidation (redox) imbalance plays a role. In this study we use the autofluorescent properties of two redox molecules, NAD(P)H and FAD, to quantify changes in their relative abundance in living lung tissue of mice with experimental lung fibrosis, and in freshly isolated cells from mouse lungs and humans with IPF. Our results identify cell population-specific intracellular redox changes in the lungs in experimental and human fibrosis. We focus particularly on redox changes within collagen producing cells, where we identified a bimodal distribution of NAD(P)H concentrations, establishing NAD(P)H^high^ and NAD(P)H^low^ sub-populations. NAD(P)H^high^ fibroblasts exhibited elevated pro-fibrotic gene expression and decreased collagenolytic protease activity relative to NAD(P)H^low^ fibroblasts. The NAD(P)H^high^ population was present in healthy lungs but expanded with time after bleomycin injury suggesting a potential role in fibrosis progression. We identified a similar increased abundance of NAD(P)H^high^ cells in freshly dissociated lungs of subjects with IPF relative to controls, and similar reductions in collagenolytic activity in this cell population. These data highlight the complexity of redox state changes in experimental and human pulmonary fibrosis and the need for selective approaches to restore redox imbalances in the fibrotic lung.

## Introduction

There is a growing consensus that a reduction-oxidation (redox) imbalance plays a crucial role in the progression of idiopathic pulmonary fibrosis (IPF) (1–4). In particular, previous work has documented a significant increase in oxidizing (e.g., reactive oxygen species) molecules and a significant decrease in reducing molecules (e.g., antioxidants (5–7)) in IPF. An imbalanced redox state is most often associated with changes in cellular metabolism but has also been shown to contribute to many of processes which propagate fibrosis including apoptosis, invasion, migration, and senescence (8–11, 4). To date, attempts to repair the redox state, primarily using antioxidants, have largely failed (12). However, redox imbalances have predominantly been studied thus far in whole lung tissue or in cultured single lung cell types, raising the possibility that complex cell-specific changes in redox state in the fibrotic lung have been overlooked.

To measure the intracellular redox state, two autofluorescent redox cofactors, NAD(P)H and FAD, are often used and can be quantified non-invasively and non-destructively using optical techniques (13–16). NAD(P)H and FAD (as electron donor and electron acceptor, respectively) are required in many intracellular chemical reactions which require electron exchange (17, 18). Often, the redox state of a cell is characterized as a ratio of the redox cofactors, calculated by 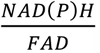, for normalization (19, 17). An increase in redox ratio is commonly associated with increased glycolysis, while a decrease in the redox ratio is associated with protein deacetylation, antioxidant formation, or poly ADP ribosylated proteins (20), all of which are known to be factors in propagating fibrosis. The goal of this study was to identify cell population-specific contributions to the redox state throughout fibrosis progression and early stages of resolution in an experimental model of pulmonary fibrosis, with comparison to fresh-dissociated cells from human fibrotic and control lungs. We used intravital imaging and flow cytometry to quantify intracellular redox state of cells in their native state or freshly isolated from normal or fibrotic tissue. We show for the first time cell population-specific changes in the redox state throughout fibrosis progression and early resolution. We found NAD(P)H level increases in fibroblasts after bleomycin administration, and NAD(P)H^high^ fibroblasts exhibit decreased intracellular collagenolytic enzyme activity. Notably, NAD(P)H^high^ fibroblasts are lost *in vitro*, preventing their further characterization using standard methods. In human samples, we found a similar increase in the NAD(P)H levels of tri-lineage negative (CD45-/CD31-/CD326-) cells which also exhibited decreased intracellular collagenolytic enzyme activity. These data suggest that a redox-shifted sub-population of fibroblasts may play a role in the progression and persistence of fibrosis.

## Materials and Methods

All experiments involving animals were performed in accordance with the guidelines and under protocols approved by the Mayo Clinic IACUC for the bleomycin induced model of fibrosis, or Ohio State University IRB for patient samples. Detailed methods and product information can be found in the supplement.

### Bleomycin Administration

Col1a1 (collagen type I alpha 1 chain)-GFP transgenic mice were a gift provided by Dr. Derek Radisky, and were generated as previously described (21). 8–12-week-old Col1α1-GFP mice were anesthetized, intubated, and bleomycin, or PBS, was administered intratracheally.

### FACS/Flow Cytometry

#### Mouse FACS isolation

Col1α1-GFP mice were anesthetized, and the lungs were perfused and harvested. We mechanically and enzymatically digested the lungs to form a single cell suspension. We lysed the red blood cells and depleted the CD45+ cell population using magnetic beads. We used flow cytometry to analyze or sort populations (endothelial, CD45-/CD31+; epithelial, CD45-/CD31-/CD326+; fibroblasts, CD45-/CD31-/CD326-/GFP+; NAD(P)H^high^ fibroblasts, CD45-/CD31-/CD326-/GFP+/NAD(P)H^high^; and NAD(P)H^low^ fibroblasts,CD45-/CD31-/CD326-/GFP+/NAD(P)H^low^) and analyzed intracellular cathepsin k activity using FlowJo. Redox ratio was calculated per cell (NAD(P)H/FAD).

#### Mouse cathepsin k flow cytometry analysis

Part of the CD45 depleted sample (described above) was added to one half of the volume of antibodies described above supplemented with MagicRed cathepsin k assay probe (1:150) on ice for 30 min. Flow cytometry was performed on a BD Fortessa x-20. Cathepsin k activity was determined using a 561 nm laser (gating strategy: Supp Fig 7). Samples were analyzed using FlowJo (v10.8, BD).

#### Human FACS/FLOW analysis

Human sample collection was performed at Ohio State University. Tissue was mechanically and enzymatically digested. Red blood cells were lysed, and samples were shipped to Mayo Clinic for further analysis. After thawing, cells were processed (CD45 depletion, FACS, FLOW) to identify cellular populations as described above.

### Longitudinal intravital imaging and image analysis

Surgical implantation of a permanent thoracic window was performed as previously described (24) with minor modifications.

For intravital imaging, mice were anesthetized and imaged using a 2-photon microscope. NAD(P)H and FAD were excited at 800 nm and collected using a 505 nm dichroic mirror and a 460-500 nm filter, for NAD(P)H, or a 520-550 nm filter, for FAD. GFP was excited at 488 nm. To analyze the images, we used a custom script in ImageJ (28).

### Primary Cell Culture

For repeated flow cytometry sorting and analysis, FACS-isolated mouse lung fibroblasts were plated in 6-well dishes (100,000 cells per well), then lifted for reanalysis weekly.

### qPCR

The mRNA of FACS-isolated cells was isolated and then converted into cDNA. qPCR of cDNA was performed using primers (Supp Table 2).

### Fibrosis evaluation

5 μm sections from FFPE lung tissues, and the sections were stained either with H&E or with Masson’s Trichrome Stain Kit.

Hydroxyproline content was measured using an acid-based hydroxyproline assay kit and read at OD 550 nm compared to a hydroxyproline standard curve.

### Statistics

All data are presented as the mean ± SEM. Statistical analysis for each figure is provided in each figure caption. Statistics were not performed on intravital imaging results due to low sample numbers.

## Results

We used flow cytometry to assess redox-related autofluorescence (16, 23) in freshly isolated cells as a primary method to analyze their *in vivo* redox state. We used Col1α1-GFP mice to assist with positive identification of definitive fibroblasts, and administered a single dose of bleomycin with tissue collected 14- or 35-days after bleomycin administration to harvest cells near the peak fibrosis, and early during the spontaneous resolution of fibrosis that occurs in young mice, respectively (29, 30).

Figure 1 shows the relative NAD(P)H, FAD, and redox ratio (NAD(P)H/FAD) for epithelial, endothelial, and tri-lineage negative (CD45-/CD326-/CD31-) cells at baseline and at days 14 and 35 after bleomycin exposure. Our findings indicate there were no significant changes in epithelial and endothelial NAD(P)H nor redox ratio levels across timepoints (Fig. 1A,B,G,H). Both endothelial and epithelial populations exhibited an increase in FAD at day 35 (Fig. 1D,E). FAD levels are closely related to those of another redox coenzyme, NAD+, which has been previously linked to fibrosis resolution (31, 32, 11). Tri-lineage negative cells, on the other hand, showed a significant decrease in NAD(P)H autofluorescence 14 days after bleomycin administration (22% decrease compared to sham), followed by a partial recovery 35 days after administration (12% increase from day 14, Fig. 1C). In this cell population FAD levels were relatively unchanged (Fig. 1F), and the overall redox ratio was reduced as result of the NAD(P)H changes (Fig. 1I). These data demonstrate differences in redox responses within sorted lung cell populations isolated after bleomycin injury, supporting cell population specific responses during fibrotic lung remodeling.

**Figure 1.**
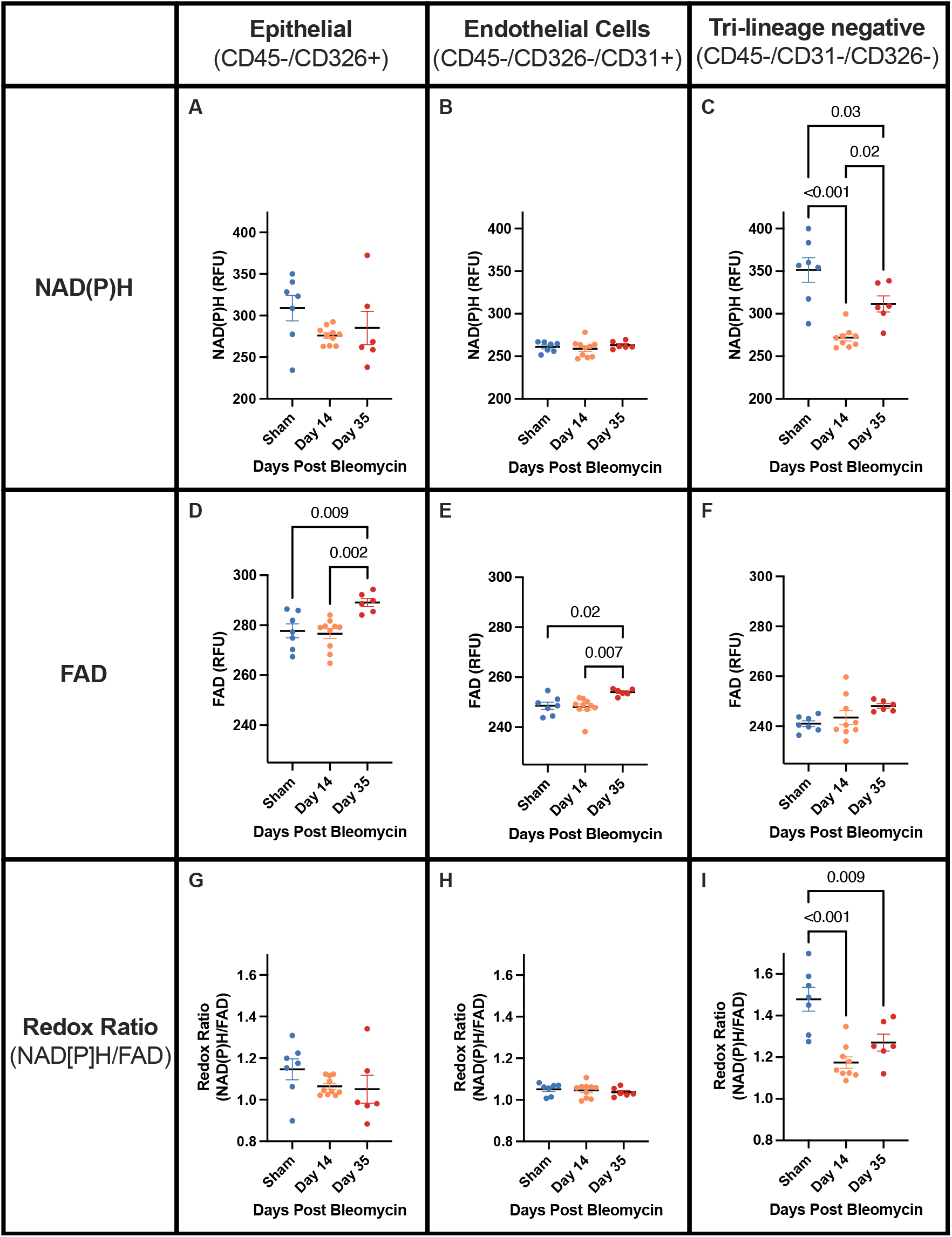
Redox changes in freshly isolated lung populations across lung fibrosis and early repair. A-C) mean NAD(P)H autofluorescence; D-F) mean FAD autofluorescence; and G-I) mean redox ratio (NAD[P]H/FAD) of A,D,G) CD45-/CD326+ epithelial cells; B,E,H) CD45-/CD326-/CD31+ endothelial cells; C,F,I) CD45^-^/CD31^-^/CD326^-^ tri-lineage-negative cells. Each point indicates the arithmetic mean value from all indicated cells from one animal. One-way ANOVA with Tukey’s post-hoc test was run for statistical analysis; n=6-10.

Potential limitations of the approach above include the time lag and disruption of native cell environments needed for lung cell isolation, both of which may alter the native redox state that exists *in vivo*. To overcome this limitation, we used a thoracic window to allow us to visualize longitudinal redox changes *in vivo* and compared tissue level *in vivo* redox changes to those observed in freshly isolated cells. Previous reports of implanted thoracic windows for longitudinal intravital imaging were limited to 14 days (24, 25). Here we sought to extend the duration to 35 days and perform long-term longitudinal studies over the course of lung injury, fibrosis, and early stages of resolution. We first tested the feasibility of thoracic window implantation in the mouse bleomycin model of lung fibrosis, and specifically evaluated whether thoracic window implantation would alter the course of lung injury, fibrosis, and repair. We then compared the responses of control and window-implanted mice to bleomycin administration at days 21 and 35, and observed similar degrees of fibrosis and resolution, as measured by hydroxyproline levels, in both groups (Supplemental figure 2).

Having validated the bleomycin model of fibrosis and repair in window-implanted mice, we then used a 2-photon microscope to measure tissue redox levels via autofluorescence at days 14, 21 and 35, spanning the time course of bleomycin-induced fibrosis and repair. For comparison, we isolated single cell suspensions of CD45-depleted cells from the same timepoints and analyzed their redox state by flow cytometry as above (timeline of experiment Fig. 2A). Both freshly isolated tissue resident cells and intravital imaging indicated a decrease in NAD(P)H levels 14 days after bleomycin administration (10% and 20% reduction, respectively) which returned to sham levels 21 days after bleomycin administration and remained stable at 35 days after bleomycin administration (Fig. 2B,C). Representative images of the NAD(P)H autofluorescence confirmed the transient reduction of NAD(P)H autofluorescent signal 14 days post bleomycin administration (Supp. Fig. 3). FAD levels did not measurably change using either *in vivo* imaging or flow cytometric analysis of lung cells (Fig. 2D,E), although more variability was observed in the intravital analysis. The redox ratio was significantly reduced at day 14 by as measured by flow cytometry (Fig. 2F), though not by intravital imaging (Fig. 2G), likely due to increased variability in the FAD autofluorescence. The general agreement between trends in intravital and dissociated cell redox changes reassured us as to the validity of the latter approach. The redox differences we observed above in cell populations, particularly the striking changes seen in the lineage negative population over time, prompted us to next examine changes in collagen-expressing fibroblast subpopulations in greater depth.

**Figure 2.**
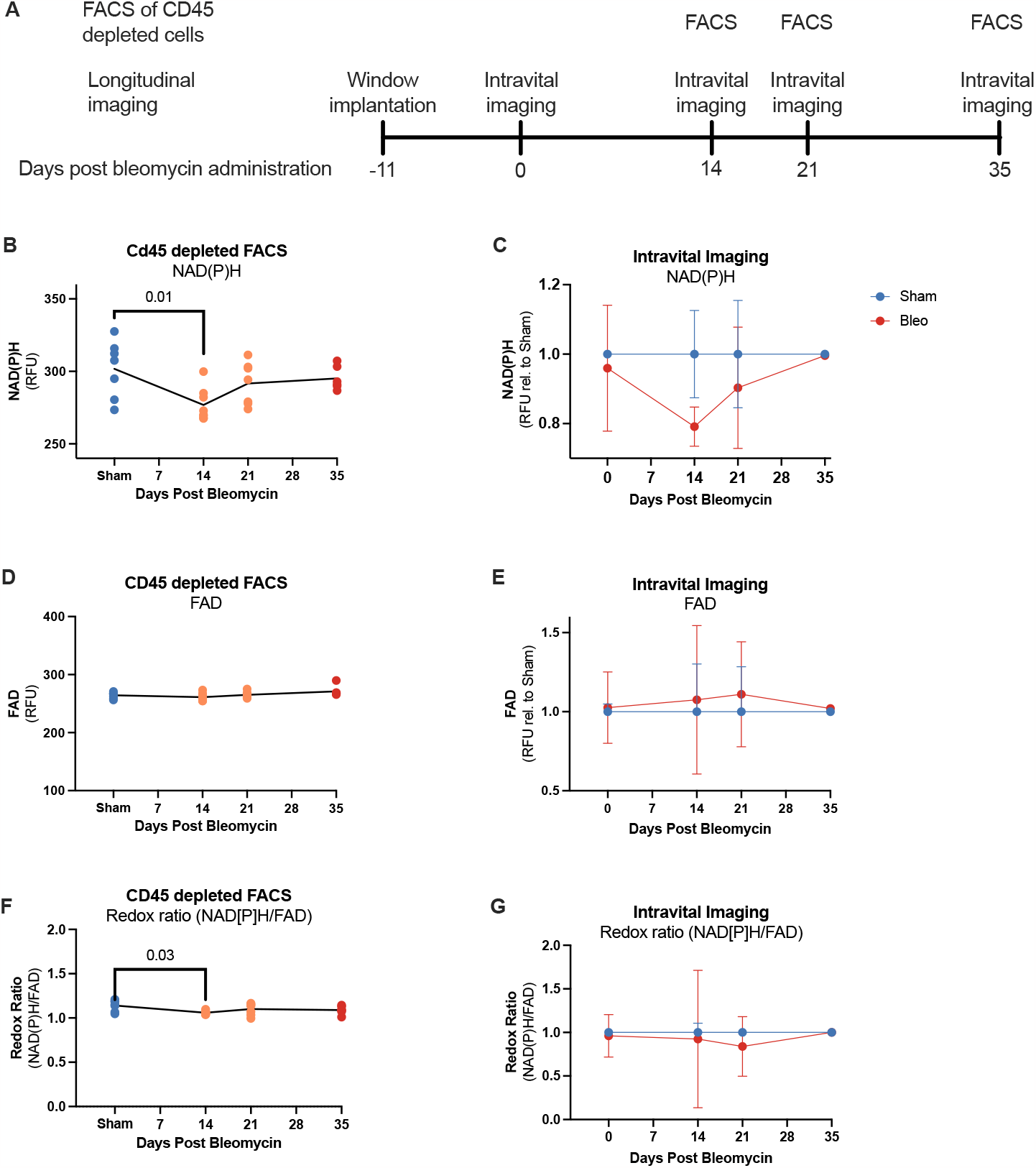
Freshly isolated cells and intravital imaging demonstrate similar NAD(P)H changes in a bleomycin model of fibrosis. A) Schematic and timeline of longitudinal intravital imaging and FACS isolation of freshly isolated cells. B) Longitudinal studies of intravital imaging show large variability in redox ratios (NAD(P)H/FAD; quantified from 2-photon images). C) Longitudinal studies of intravital imaging show large variability in FAD levels in after bleomycin administration quantified from 2-photon images. D) Longitudinal studies of intravital imaging show decreased NAD(P)H levels 14 days after bleomycin administration, quantified from 2-photon images. For B-D, each point is the arithmetic mean of all animals imaged. No statistical analysis was performed for intravital imaging; n=1 per group at Day 35, n=4 at all other time points; intravital imaging studies were longitudinal. E) The redox ratio (NAD[P]H/FAD) of resident lung cells is significantly decreased 14 days after bleomycin administration. F) FAD is relatively unchanged at various timepoints after bleomycin administration. Points indicate measured autofluorescence of FAD from freshly isolated resident lung cells taken from independent samples. G) NAD(P)H is significantly decreased in resident lung cells 14 days post bleomycin administration. Points indicate measured autofluorescence of NAD(P)H from freshly isolated lung resident cells taken from independent samples. For E-G, each point is the arithmetic mean of all CD45-cells isolated from a single animal. One-way ANOVA with Tukey’s post-hoc test run for statistical analysis; n=6-7.

We thus repeated the flow-based assessment of redox state in the Col1a1-GFP+ fraction of tri-lineage-negative cells. Strikingly, these cells differed from the parent tri-lineage-negative population by maintaining NAD(P)H levels 14 days after bleomycin administration, but then significantly increasing NAD(P)H levels 35 days after bleomycin administration (6% increase compared to sham; Fig. 3A). The results in Figure 3A show the average NAD(P)H levels across animals, but we also noted striking variation in NAD(P)H levels within this cell population. When we plotted the distribution of NAD(P)H levels in the Col1a1-GFP+ fraction of tri-lineage-negative cells, we identified a bimodal distribution of NAD(P)H levels (Fig. 3B). A similar bimodal distribution was noted in the parent tri-lineage-negative population, but not in endothelial or epithelial populations (Supp. Fig. 4), establishing the existence of unique NAD(P)H^low^ and NAD(P)H^high^ fibroblast subpopulations within the lung. Interestingly, the fraction of NAD(P)H^high^ fibroblasts increased after bleomycin administration, becoming significantly increased 35 days after bleomycin administration (67% increase over sham; Fig. 3C). The bimodal distribution of NAD(P)H suggested that these may constitute redox-specific functional sub-populations of fibroblasts.

**Figure 3.**
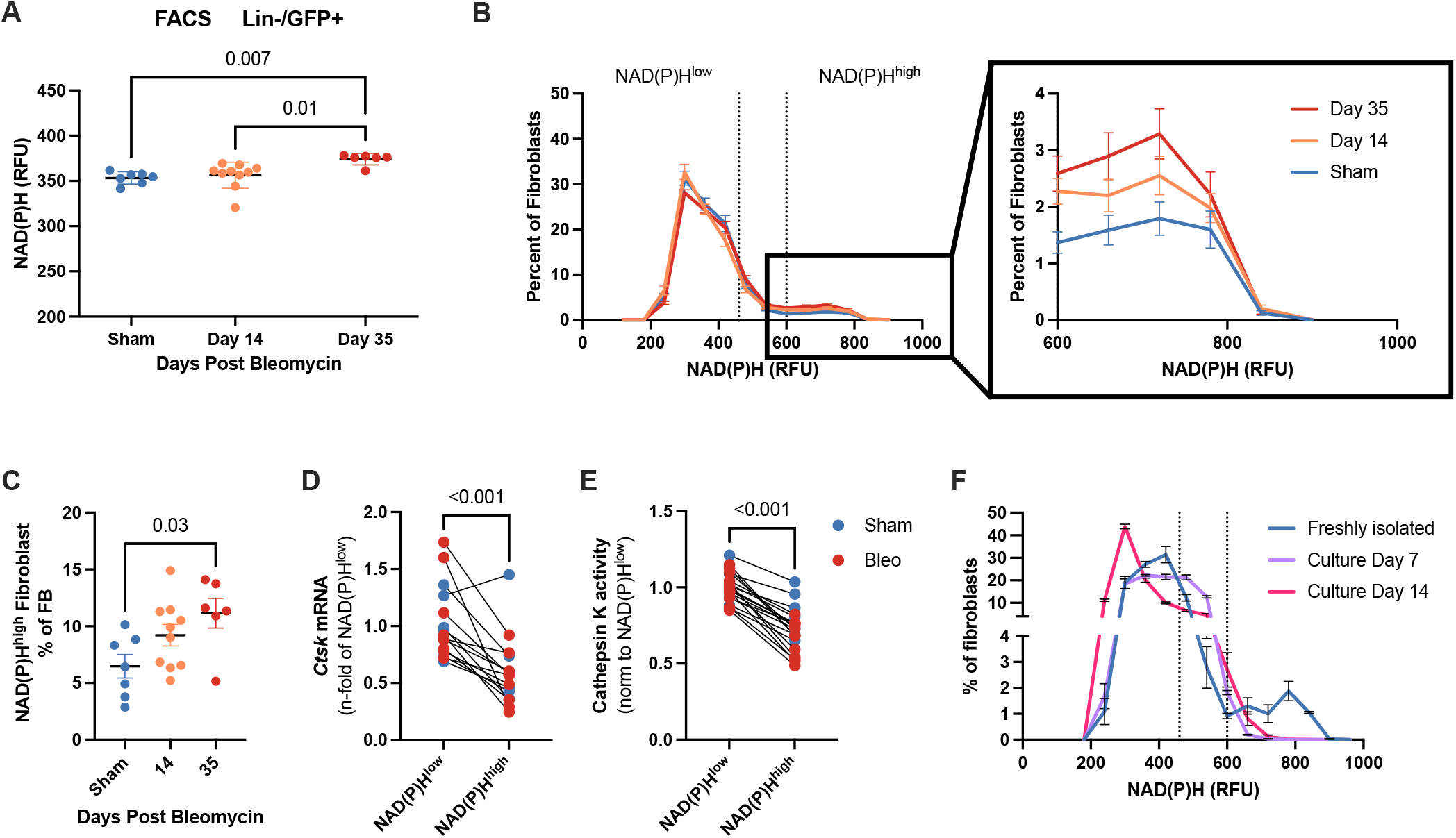
Fibroblasts exhibit redox specific sub-populations with decreased intracellular cathepsin k activity in NAD(P)H^high^ fibroblasts. A) In a bleomycin model of fibrosis NAD(P)H^high^ fibroblasts increase 35 days after bleomycin administration. Each point indicates the combined arithmetic mean of all Lin-/GFP+ (CD45-/CD326-/CD31-/GFP+) fibroblasts from one animal; n=6-10. B) Longitudinal studies of intravital imaging show slight decreases in NAD(P)H levels 14 days after bleomycin administration, increasing to above sham level 35 days after bleomycin administration. Values quantified from 2-photon images. N=4 for each group at day 0 and day 14, n=1 for day 35. Statistical analyses were not performed on these samples. C) Fibroblasts have a bimodal NAD(P)H distribution with increased NAD(P)H^high^ fibroblasts 35 days after bleomycin administration. D) NAD(P)H^high^ fibroblasts are significantly increased 35 days after bleomycin administration. Each point indicates the combined arithmetic mean of all Lin-GFP+ fibroblasts from one animal; n=6-10. E) NAD(P)H^high^ fibroblasts have decreased expression of *Ctsk*. Each point indicates the n-fold change (2^) −(ΔΔ*cq*)^) relative to the average of all NAD(P)H^low^ fibroblasts from vehicle control animals. n=15. F) Freshly isolated NAD(P)H^high^ fibroblasts have decreased intracellular cathepsin K activity quantified through flow cytometry. Each point indicates the combined arithmetic mean of all Lin-GFP+ fibroblasts from one animal; n=20. One-way ANOVA with Tukey’s post-hoc test was run for statistical analysis. G) NAD(P)H is decreased in cell culture. NAD(P)H^high^ cells are lost within 7 days of cell culture. n=5. H) FAD increases in within 7 days of cell culture. n=5.

To identify phenotypic differences between these redox-distinct populations, we analyzed cells from multiple timepoints after bleomycin administration, seeking to test whether a distinct NAD(P)H-based phenotype exists independent from cell changes that occur in fibrosis. Based on recent work from our group implicating fibroblast expression of cathepsin K as potentially important in the resolution of fibrosis at later stages after bleomycin administration (33), we analyzed *Ctsk* transcript levels (encoding cathepsin K) and noted significantly lower levels in NAD(P)H^high^ compared to NAD(P)H^low^ fibroblasts; these differences were independent of whether the cells were sorted from control or bleomycin-injured mice (Fig 3D). Other mRNA transcript levels suggested a pro-fibrotic phenotype of NAD(P)H^high^ fibroblasts, including increased NADPH oxidase 4 (*Nox4*) and decreased cyclooxygenase 2 (*Ptgs2*), while ECM genes collagen I alpha I (*Col1a1*) and fibronectin (*Fn1*), and collagenolytic matrix metalloprotease 14 (*Mmp14*) were unchanged (Supp. Fig. 5). These results suggest both subpopulations to be fibroblasts, with NAD(P)H^high^ fibroblasts exhibiting redox-specific differences in key signaling programs.

To build on the observed differences in *Ctsk* expression, we next sought to directly assess activity of cathepsin K protein. Importantly, cathepsin K can promote collagen resorption (33) but this collagenolytic activity can also be post-translationally modified (34, 35), requiring direct measurement of activity. To measure intracellular cathepsin K activity, we used a cell-permeable cathepsin k probe (MagicRed) which localizes to the lysosomes and fluoresces when cleaved by cathepsin K. We exposed freshly isolated cells to MagicRed and used flow cytometry to analyze cathepsin K activity in NAD(P)H^high^ and NAD(P)H^low^ populations. Cathepsin k activity was significantly lower (20% reduction in fluorescence levels) in NAD(P)H^high^ compared to NAD(P)H^low^ fibroblasts (Fig. 3E). These data suggest that NAD(P)H^high^ fibroblasts may have lower capacity to degrade type I collagen, with potentially important implications for fibroblast subpopulations that promote fibrosis progression versus resolution. Further confirmation of functional differences in NAD(P)H-specific populations would benefit from *in vitro* studies.

However, when we cultured a mixed population of NAD(P)H^high^ and NAD(P)H^low^ fibroblasts, the NAD(P)H^high^ subpopulation disappeared within seven days, leaving further functional study to await development of culture methods capable of maintaining this fibroblast subset ex vivo (Fig. 3G).

Despite our inability to dissect the biology of NAD(P)H^high^ fibroblasts *in vitro*, we were intrigued by their appearance in the late stages of the bleomycin model and sought to assess whether similar redox subpopulations of fibroblasts may exist in the fibrotic human lung. We repeated our FACS and flow cytometry-based redox characterization of cells dissociated from human control and IPF lungs. In contrast to the bleomycin model, we observed more differences in NAD(P)H levels and fewer differences in FAD levels between control and fibrotic lung cell populations (Fig. 4). Notably, IPF-derived tri-lineage-negative cells, commonly found to be predominantly fibroblasts in human lungs (36, 37), exhibited higher NAD(P)H levels in aggregate than the same cells from control lungs (Fig. 4C), similar to the Col1α1-GFP+ fibroblasts at day 35 after bleomycin injury.

**Figure 4.**
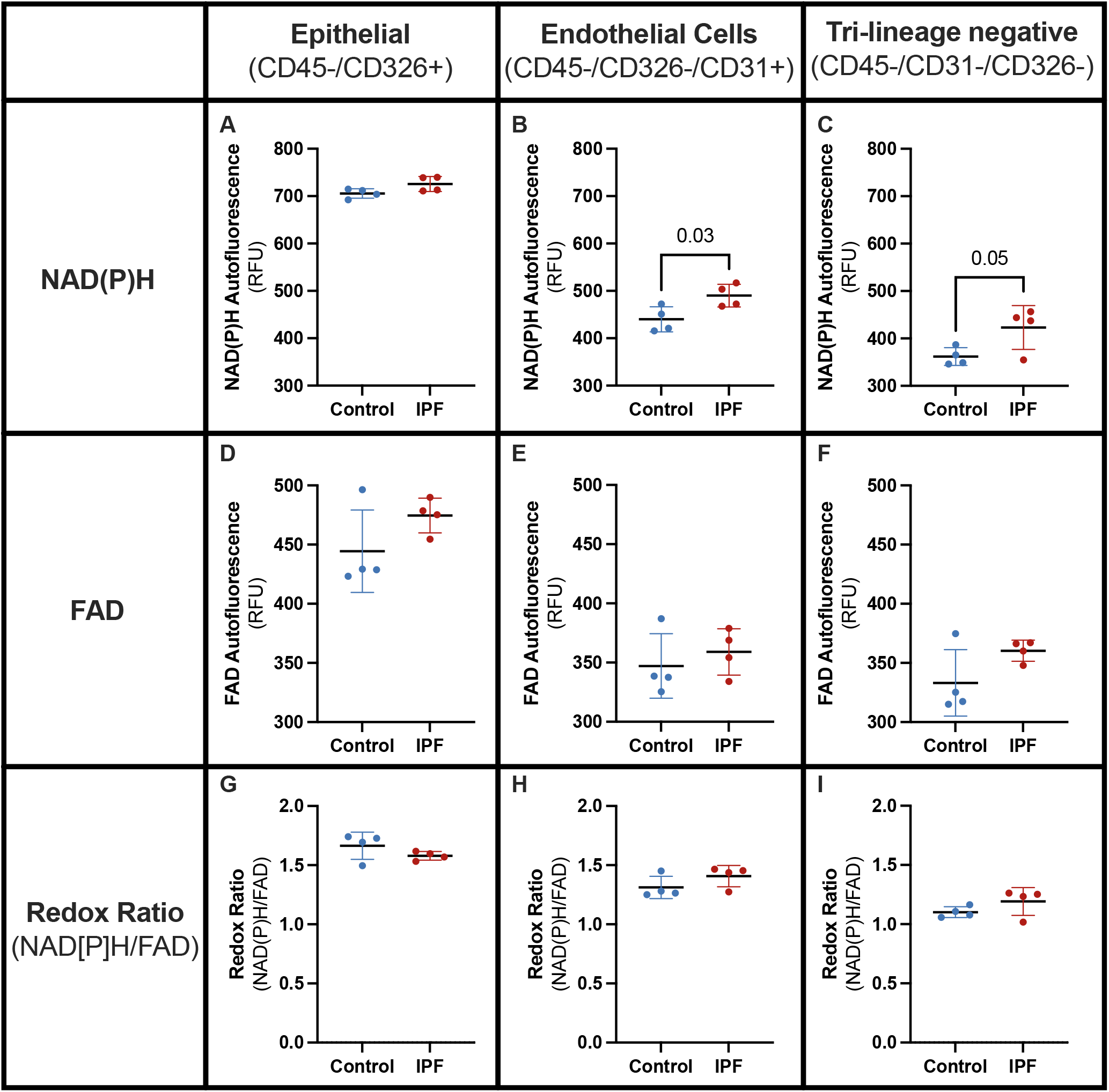
NAD(P)H increases in endothelial cells and mesenchymal cells isolated from human IPF, while FAD and redox ratios remain relatively unchanged. A-C) mean NAD(P)H autofluorescence; D-F) mean FAD autofluorescence; and G-I) mean redox ratio (NAD[P]H/FAD) of A,D,G) CD45-/CD326+ epithelial cells; B,E,H) CD45-/CD326-/CD31+ endothelial cells; C,F,I) CD45^-^/CD31^-^/CD326^-^mesenchymal cells. Each point indicates the arithmetic mean value from all indicated cells from one patient. One-way ANOVA with Tukey’s post-hoc test was run for statistical analysis; n=4.

To identify if the tri-lineage-negative human lung cell subpopulation exhibited similar changes in the distribution of NAD(P)H levels, we further investigated this population by flow cytometry as we did for mouse lung cells. Using the (CD45-/CD326-/CD31-) populations, we did not observe as distinct a bimodal NAD(P)H distribution, but we did observe a dramatic increase in the width of the NAD(P)H distribution with a right-shift of NAD(P)H levels in IPF versus control lung cells (Fig. 5A). This right-shift of NAD(P)H levels corresponded to a near doubling of NAD(P)H^high^ cells (85% increase) in IPF versus control cells, though with considerable variability across human IPF samples (Fig. 5B). Finally, we considered whether the NAD(P)H^high^ cells might exhibit a functional phenotype similar to our observations in mice, irrespective of IPF diagnosis. Consistent with our results from the NAD(P)H-associated phenotype of mice, NAD(P)H^high^ cells exhibited a lower expression of *CTSK* (Fig. 5C) and a small but significant reduction in cathepsin k activity (9% decrease in fluorescence; Fig. 5D). Together these results suggest that redox-associated fibroblast subpopulations exist in the human lung, and may exhibit phenotypic traits that contribute to the persistence of fibrosis and inability to degrade collagen I.

**Figure 5.**
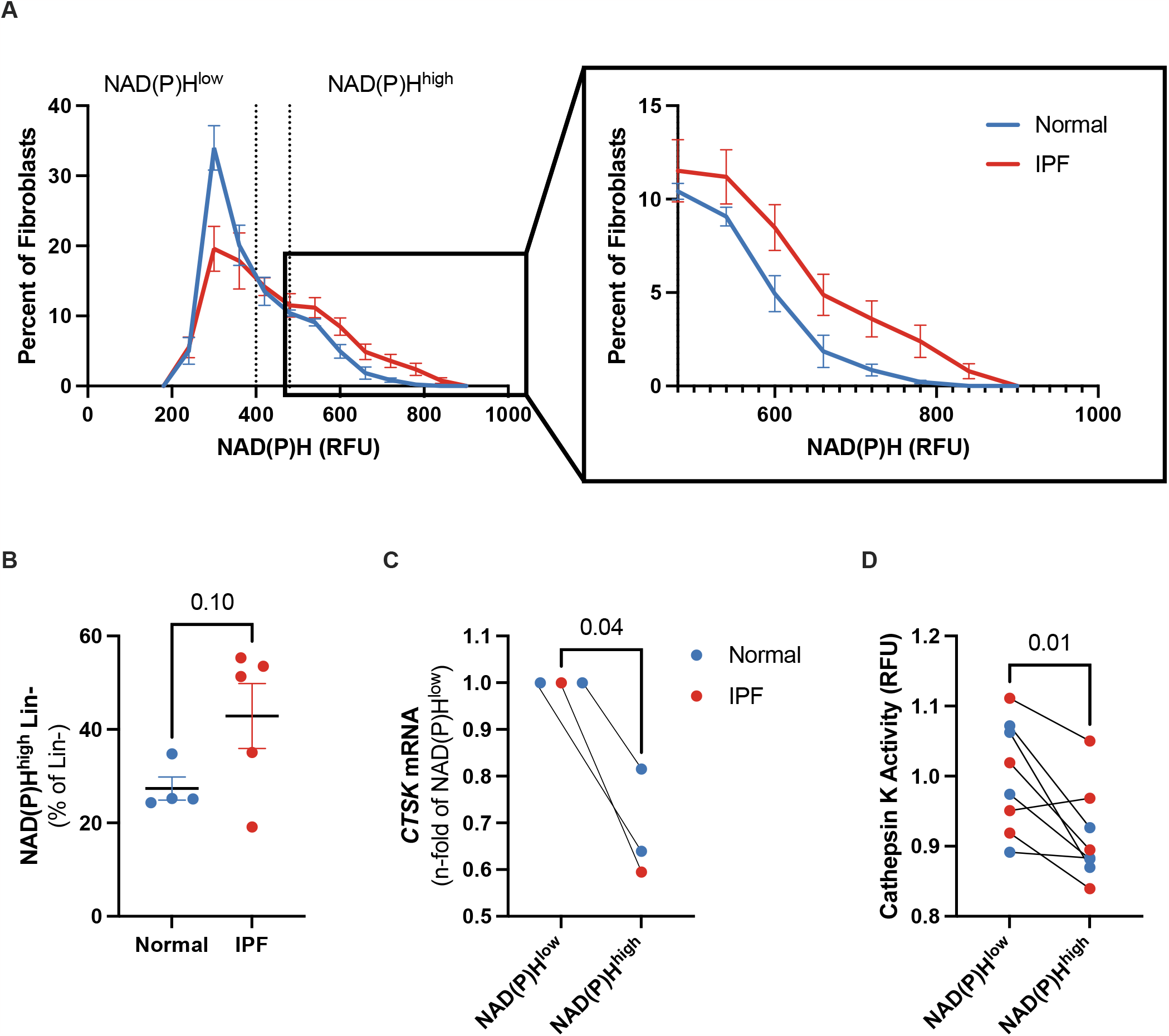
In IPF-derived mesenchymal cells, NAD(P)H^high^ cells have decreased cathepsin k activity. NAD(P)H is shifted toward higher NAD(P)H levels. A) Although the bimodal NAD(P)H distribution disappears, a right shift of increased NAD(P)H exists in IPF samples B) NAD(P)H-hi cells nearly double in IPF. n=4-5 C) *CTSK* mRNA expression is reduced in NAD(P)H^high^ cells. n=3. D) Cathepsin K activity is reduced in NAD(P)H^high^ cells. Each point indicates the combined arithmetic mean of all Lin-cells from one patient; n=8. In each plot, a t-test was used for statistical analyses, unpaired when comparing normal to IPF (B) and paired when comparing NAD(P)H^high^ to NAD(P)H^low^(C,D).

## Discussion

Our results characterize for the first-time population-specific intracellular redox changes throughout fibrosis progression and repair. We focused particular attention on freshly isolated collagen producing fibroblasts during fibrosis and resolution given their central role in matrix deposition and potential roles in matrix resorption. In fibroblasts, we identified little change in FAD but significant changes in NAD(P)H. Interestingly, we found that fibroblasts have a bimodal distribution of NAD(P)H, establishing an NAD(P)H^high^ population and an NAD(P)H^low^ population. We began characterizing the differences in the NAD(P)H^high^ and NAD(P)H^low^ fibroblasts and found gene expression differences and collagenolytic protein activity changes, with NAD(P)H^high^ fibroblasts exhibiting increased pro-fibrotic signals and reduced collagenolytic and anti-fibrotic signals. The NAD(P)H^high^ population increased with time after bleomycin and may identify a sub-population of fibroblasts which play a role in fibrosis progression and prevent resolution. We validated these observations in cells isolated from human IPF patients and found similar changes with reduced collagenolytic expression and activity in NAD(P)H^high^ cells.

In the bleomycin model of fibrosis, intracellular redox changes showed decreases in NAD(P)H during fibrosis progression, while FAD increased during resolution. Combining individual cell redox states to reconstruct tissue-level redox changes largely aligned, albeit with a small number of individuals, with *in vivo* 2-photon imaging of NAD(P)H levels. The reduction in NAD(P)H levels 14 days post bleomycin administration could indicate an increase in glycolysis, known to be involved in the progression of fibrosis (38, 39). Similarly, the increases in FAD levels during the resolution phase could represent not only a restoration in metabolic oxidative phosphorylation activity, but also an increased production of NAD+ to increase antioxidant functionality through sirtuins, both of which have been shown to be important in resolving models of fibrosis (38, 40, 41). Current literature suggests restoring NAD+ levels to resolve experimental fibrosis (11, 31, 32), and our work supports those changes through the significant increases in FAD (as a surrogate marker for NAD+; (13, 42, 39)) during resolution timepoints. While the major cell populations generally aligned in redox shifts after bleomycin administration, collagen producing GFP+ fibroblasts did not.

Fibroblast NAD(P)H increased with time after bleomycin administration, which presented as an increasing NAD(P)H^high^ sub-population. While other studies have identified cellular changes based on redox levels *in vitro* (10, 43), we believe this is the first report of the redox state of freshly isolated fibroblasts. We and others have previously shown cathepsin k to be an important regulator of lung collagen and a possible therapeutic target for resolving pulmonary fibrosis (44, 45, 33), and we show here NAD(P)H^high^ fibroblasts have reduced cathepsin k activity, suggesting the NAD(P)H^high^ population is a pro-fibrotic fibroblast population. Many groups have identified *Nox4* and *Ptgs2* as pro-fibrotic and pro-resolving indicators, respectively (46–49), and we identified significant changes in expression levels of those genes in NAD(P)H^high^ fibroblasts, further suggesting the NAD(P)H^high^ population is a pro-fibrotic fibroblast population. We identified similar changes in IPF-derived tri-lineage-negative cells, with increases in NAD(P)H levels with disease and decreased collagenolytic activity in NAD(P)H^high^ cells. Further characterization of this redox-imbalanced subpopulation is needed to understand the possible role of these fibroblasts in both health and diseased states; however, when we attempted to expand fibroblasts *in vitro*, we found substantial changes in the redox state as early as 7 days after isolation. To further study this population of cells will require *in vitro* techniques which retain the *in vivo* redox state, and lineage tracing studies to determine if this population is a transient population of fibroblasts. These results highlight a redox-imbalanced fibroblast sub-population, which may play a role in propagating pulmonary fibrosis and needs further characterization.

In conclusion, we have identified a shift in the redox state of cell populations throughout fibrosis, with a unique fibroblast sub-population expanding after bleomycin administration. Early characterization of these NAD(P)H^high^ fibroblasts indicates a pro-fibrotic program which may contribute to the non-resolving nature of IPF. It is important to note that these studies were performed in young animals, and further work will be needed in an aged model of fibrosis to assess overlap or distinct changes, as aging has been shown to decrease NAD+ (31, 50, 51) and an aged model of fibrosis might mimic the changes seen in the IPF-derived cell redox state better than the young bleomycin model. The observations presented here further our understanding of the redox state of freshly isolated cell populations from the fibrotic lung and should prompt further detailed evaluation of redox states in additional subpopulations of cells present in during lung fibrosis and repair. More broadly, our results highlight the need to study the role of redox in fibrosis in greater detail and underscore the complexity of targeting redox globally, raising the likelihood that cell-specific approaches may be needed to restore redox imbalances in specific lung cell populations.

## Acknowledgements

The authors would like to acknowledge the staff at the Mayo Clinic Microscopy and Cell Analysis Core for their assistance and expertise. We would also like to thank Julian A. Chamucero for assistance with the human samples. Funding support was provided by the National Institutes of Health grants HL105355, HL158018, and HL153026 (to P.A.L.), HL092961, HL166187 and HL152967 (to D.J.T.).

## Author Contributions

PAL and DJT designed the experiments. PAL, JAM, NC, AYG, VP, and JAC performed the experiments. PAL, GBS, ACR, MR and DJT analyzed acquired data. PAL and DJT. wrote the first draft of the manuscript. All authors contributed to editing the manuscript.

## Declarations of Interests

The authors declare no competing interests.

## Notes

### Competing Interest Statement

The authors have declared no competing interest.

