## Supplemental data and methods for "A redox-shifted fibroblast subpopulation emerges in the fibrotic lung"

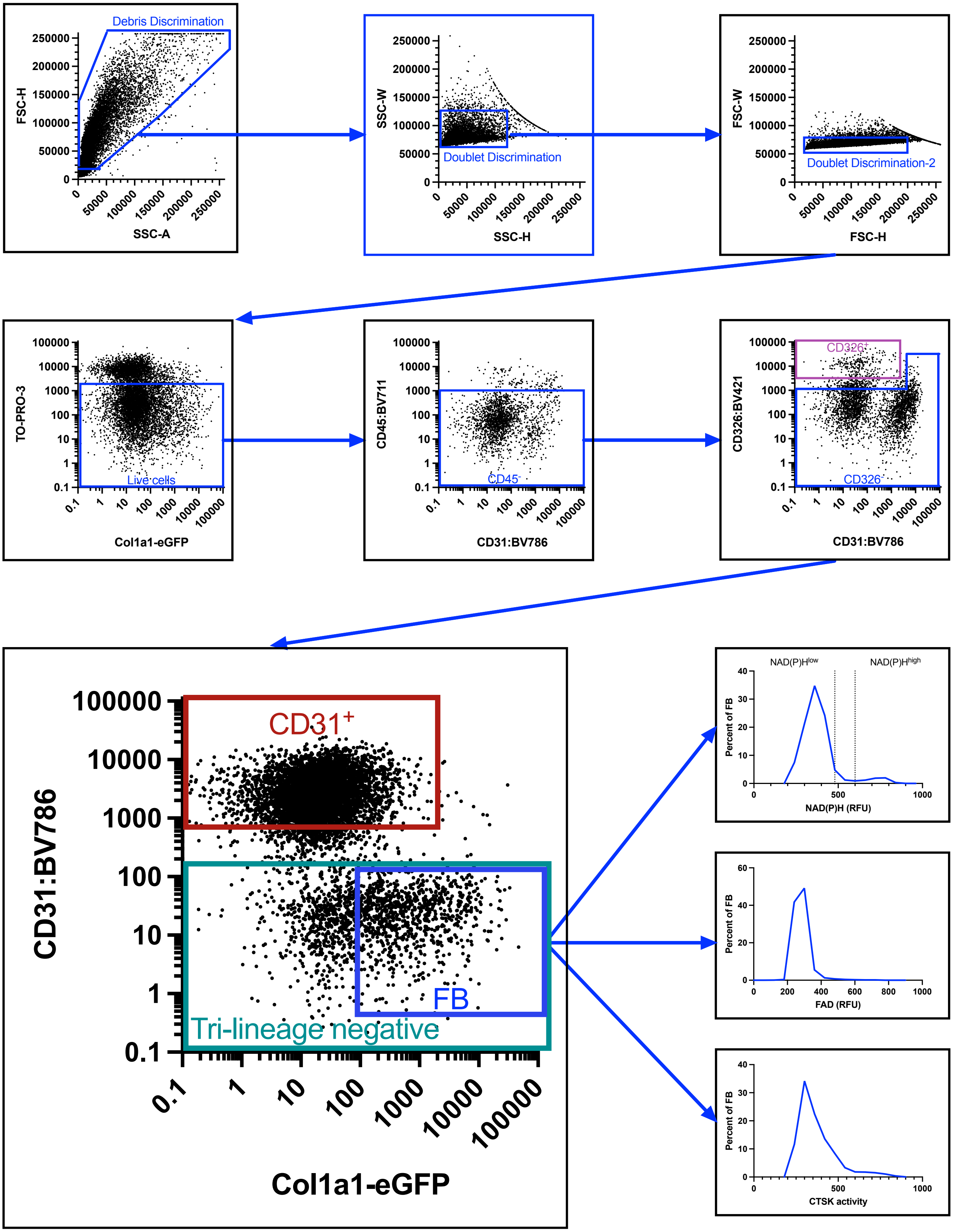

**Supp.**

**Fig. 1. Strategy for FACS-isolation/FLOW-analysis.** Lung cell suspension was depleted of CD45 cells using magnetic beads, then stained with antibodies to isolate NAD(P)H$^{high}$ and NAD(P)H$^{low}$ cells from CD45-/CD326-/CD31-/GFP+ fibroblasts.

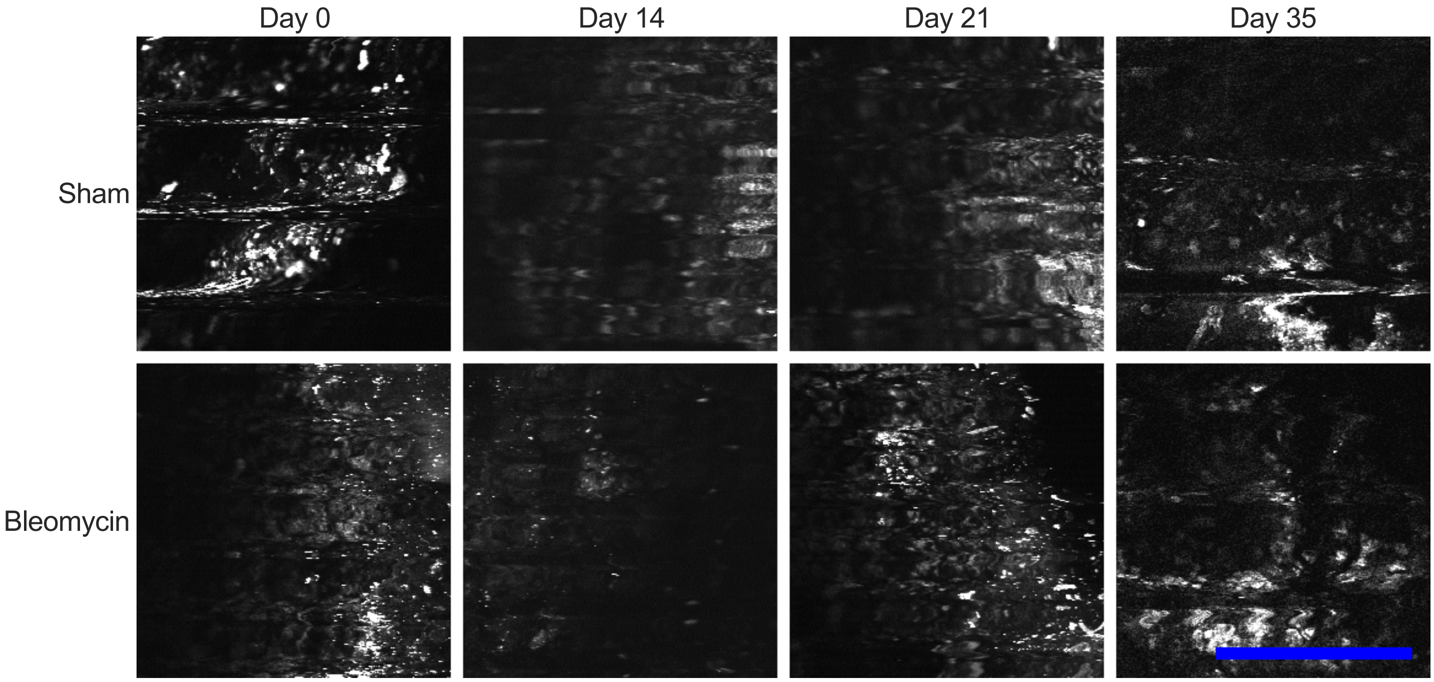

**Supp. Fig. 2. Representative images taken from intravital 2-photon imaging show decreased NAD(P)H autofluorescence 14 days after bleomycin administration.**

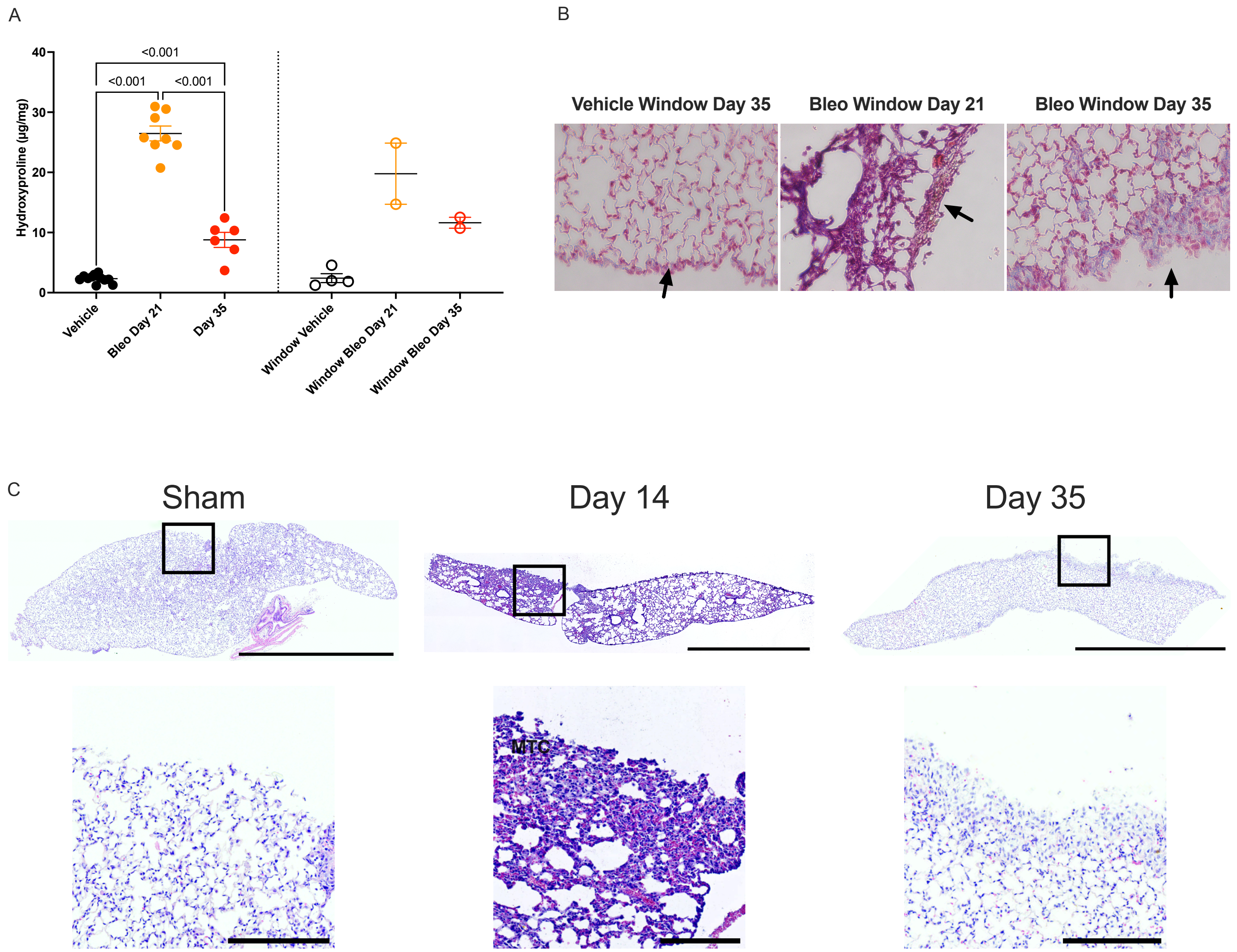

**Supp. Fig. 3. Thoracic window implantation allows for fibrosis progression and resolution.**
A) Hydroxyproline of right lungs isolated from non-window and window-implanted mice showing similar changes in total lung content. C) Masson's trichrome staining of window implanted lungs. C) Whole lung H&E of lungs isolated after window implantation and bleomycin administration. Boxes indicate lung region attached to the thoracic window, which are blown up below.

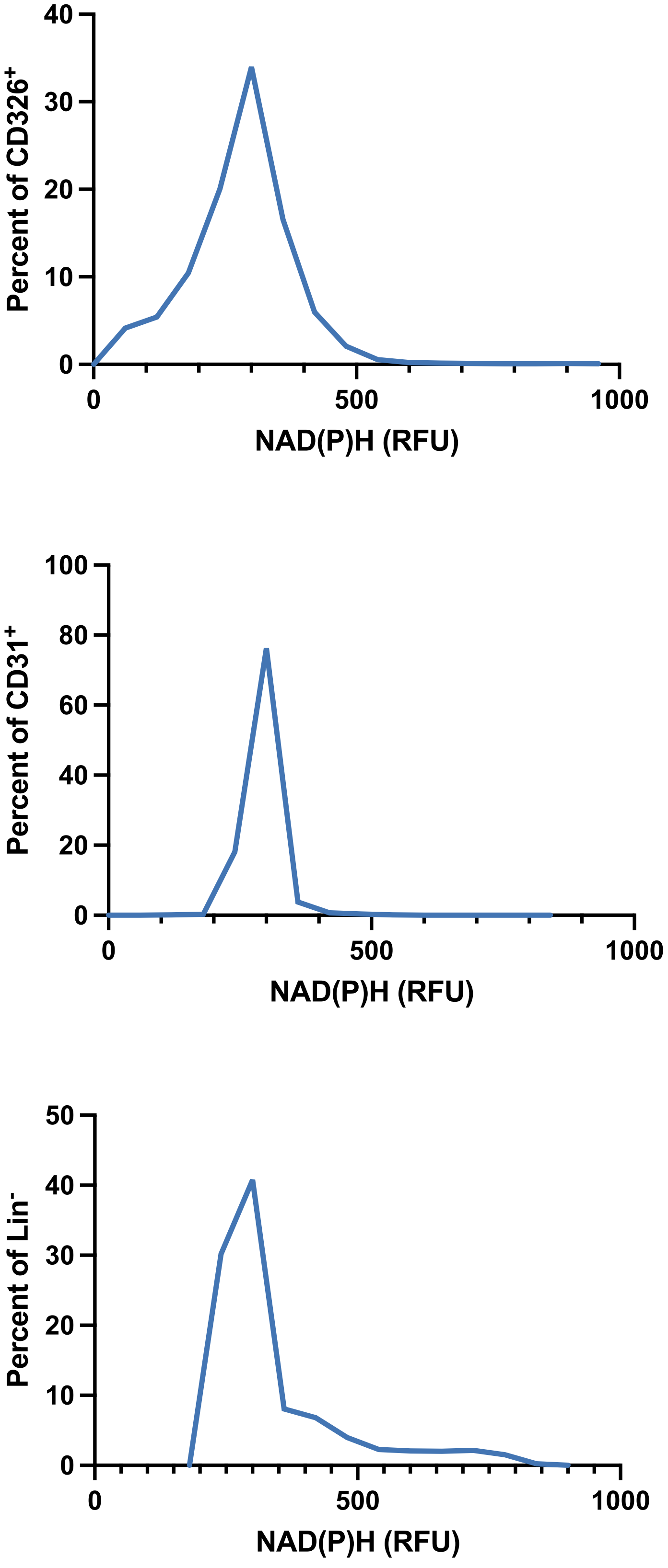

**Supp. Fig. 4. Lin- cells, but not endothelial or epithelial cells, display bimodal NAD(P)H levels.**

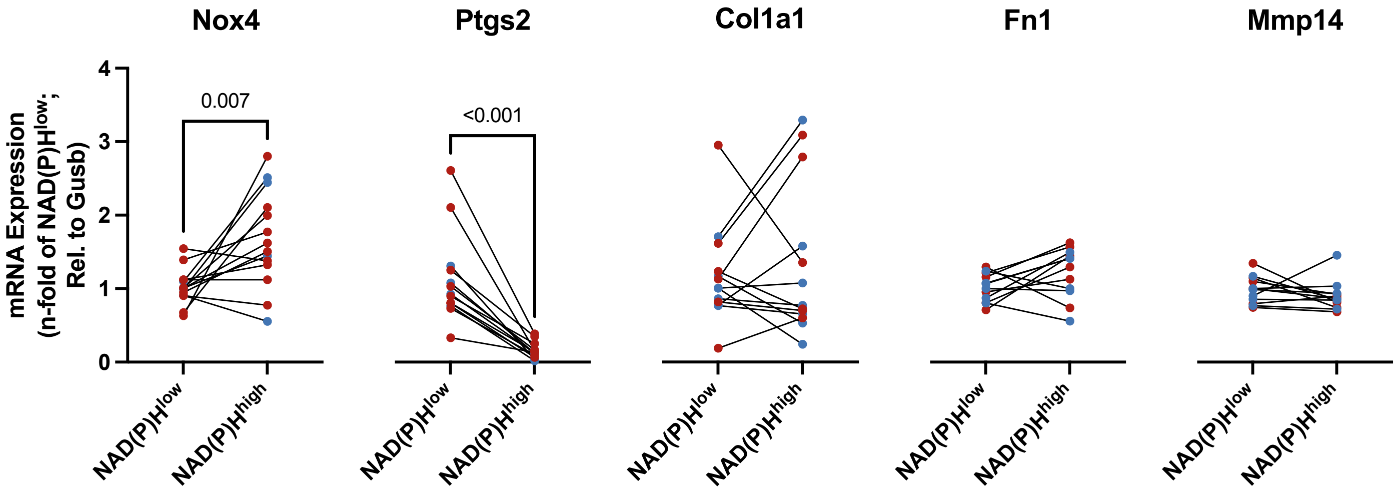

**Supp. Fig. 5. NAD(P)H$^{high}$ fibroblasts have similar decreases in identified genes, while ECM genes remain similar.** In NAD(P)H$^{high}$ fibroblasts isolated from control and bleomycin-treated mice have similar gene expression profiles regardless of treatment condition.

**Materials and Methods**
All experiments involving animals were performed in accordance with the guidelines and under a protocol approved by the Mayo Clinic Institutional Animal Care and Use Committee. All experiments involving patient samples were performed under a protocol approved by the Ohio State University Institutional Review Board. Detailed product information can be found in the supplemental information (supp table 1).

Bleomycin Administration
Col1a1 (collagen type I alpha 1 chain)-GFP transgenic mice were a gift provided by Dr. Derek Radisky (Mayo Clinic, Jacksonville, FL), and were generated as previously described (1). 8–12-week-old Col1α1-GFP mice were anesthetized with ketamine/dexdomitor solution (100 mg/kg and 1 mg/kg, respectively) injected intraperitoneally. Mice were intubated using a 20g catheter (BD, angiocath 26742) and tracheal placement was confirmed using a PBS bubble (2). We then administered bleomycin (0.7 U/kg; Hospira Inc.) in 50 µl of PBS using a MicroSprayer® (model IA-1C) Aerosolizer (FMJ-250) from Penn-Century (Wyndmoor, PA). We moved the mice to a heating pad for recovery and then reversed anesthesia using Antisedan (10 mg/kg) injected intramuscularly.

FACS/Flow Cytometry
*Mouse FACS isolation*
Col1α1-GFP mice were anesthetized using ketamine/dexdomitor solution (100 mg/kg and 1 mg/kg, respectively) injected intraperitoneally. We perfused the right ventricle with ice-cold PBS supplemented with EDTA (2 mM final concentration) to flush blood content from the lung. We then harvested the lungs and minced them in 10 cm Petri dishes. The minced tissue was moved into digestive solution (DMEM, 0.1 mg/ml Liberase TM, and 100 U/ml DNase I; final concentrations) and incubated on a tube rotator at 37°C for 45 min. We inactivated the digestive enzymes with 1x DMEM containing 10% FBS. We filtered the cell and tissue suspension using a 40-µm filter and then centrifuged the suspension. We resuspended the cell pellet in red blood cell lysis buffer for 90 s and then diluted in 3x volume of PBS. We centrifuged the cell suspension again and resuspended in 100 µl autoMACS buffer containing 10 µl CD45 MicroBeads and incubated at 4°C for 15 minutes. We depleted the CD45+ population using LS magnetic columns according to manufacturer's directions. We centrifuged the cell suspension again and resuspended the cell pellet in 100 µl autoMACS buffer. We took 10% of the suspension for cathepsin k flow cytometry (detailed below). To the remaining 90%, we added TO-PRO-3 iodide (1 µM final concentration), anti-mouse CD326-BV421 (1.36 µg/sample), anti-mouse CD45-BV711 (0.56 µg/sample), and anti-mouse CD31-BV785 (0.24 µg/sample) to CD45-depleted freshly dissociated cells to identify major cell populations. We brought the total volume up to 200 µl with autoMACS and incubated on ice for 30 min. FACS isolation/analysis was performed on a BD Aria II (gating strategy: Supp Fig. 7). Autofluorescence was recorded for NAD(P)H (ex 355B nm; filter 450/50) and FAD (ex. 407A; filter 525/50) (3). We collected the following cell populations: endothelial (CD45-/CD31+), epithelial (CD45-/CD31-/CD326+), fibroblasts (CD45-/CD31-/CD326-/GFP+), NAD(P)H$^{high}$ fibroblasts (CD45-/CD31-/CD326-/GFP+/NAD(P)H$^{high}$), and NAD(P)H$^{low}$ fibroblasts (CD45-/CD31-/CD326-/GFP+/NAD(P)H$^{low}$). Samples were analyzed using FlowJo (v10.8, BD). Values presented were exported from FlowJo as compensated channel values. Redox ratio was calculated per cell (NAD(P)H/FAD).

*Mouse cathepsin k flow cytometry analysis*
10% of the CD45 depleted sample (described above) was added to one half of the volume of antibodies described above supplemented with MagicRed cathepsin k assay probe (1:150) on ice for 30 min. Flow cytometry was performed on a BD Fortessa x-20. Cathepsin k activity was determined using a 561 nm laser (gating strategy: Supp Fig 7). Samples were analyzed using FlowJo (v10.8, BD).

*Human FACS/FLOW analysis*
Human sample collection was performed at Ohio State University. All protocols were approved by the Institutional Review Board and the Committee for Oversight of Research at the Ohio State University. Signed informed consent forms were collected before organ procurement. Once tissue was collected, it was minced and digested with Liberase DL (1 mg/mL, Roche) in a rocker incubator (30 min, 37°C). Then DNase I (100 U/mL)

was added and incubated (30 min, 37°C). We added FBS to a final concentration of 10%, filtered (300 µm, 100 µm, 70 µm, 40 µm), and then centrifuged (500g, 7 minutes) to pellet cells. We resuspended the cell pellet in Red Blood Cell Lysis buffer and incubated (4°C, 7 minutes). PBS+10% FBS was added to stop the lysis and the samples were centrifuged (500g, 7 minutes), resuspended in PBS and refiltered (40 µm). Cells (5x10^6 cells/ml) were then frozen in 50% DMEM, 40% FBS, 10% DMSO using isopropanol for controlled cooling before shipping. After thawing, cells were processed (CD45 depletion, FACS, FLOW) as described above using TO-PRO-3 iodide (1 µM final concentration), anti-human CD326-BV421 (5 µl/ million cells), anti-human CD45-BV711 (5 µl/ million cells), and anti-human CD31-BV785 (5 µl/ million cells) to identify the same cellular populations as described above.

Longitudinal intravital imaging and image analysis

Surgical implantation of a permanent thoracic window was performed as previously described (4) with minor modifications. Briefly, 10-week-old Col1α1-GFP FVB mice were given analgesics beginning 3 days prior to the surgical procedure. On the day of the procedure, we anesthetized mice with ketamine/dexdomitor solution (100 mg/kg and 1 mg/kg, respectively) injected intraperitoneally. Then mice were intubated using a 20 ga I.V. catheter and tracheal placement was confirmed using a PBS bubble (2). The catheter was tied in place and the mouse was placed on a ventilator (300 breathes/minute, 200 µl/breath) and ketamine/dexdomitor anesthesia was reversed with Antisedan (10 mg/kg) and replaced with 2% isoflurane in 100% oxygen. The hair around the surgical site was removed and a sterile field was established on top of the mice and surrounding areas. The soft tissue was removed in a 10 mm circle and then the thoracic wall was removed in a 5 mm circle. 5-0 vicryl suture was sutured to the thoracic wall and a stainless-steel frame was inserted in the space. The suture was tightened to secure the frame in place and then a 5 mm coverslip was inserted in the middle of the frame and adhered to the lung tissue. The skin was sutured under the ledge of the frame using 3-0 vicryl and the mouse was removed from anesthesia to recover.

Prior reports have indicated little to no inflammation from the window implantation procedure (4, 5), but because we administered anti-inflammatory NSAIDs, which are known to interfere with the bleomycin model of fibrosis (6, 7), as part of the thoracic window implantation procedure, we waited 11 days after window implantation before administering bleomycin.

At the indicated timepoints, mice were given ketamine/dexdomitor (75 mg/kg and 1 mg/kg, respectively) anesthesia, injected intraperitoneally. Mice were placed on a heated pad on top of the microscope stage of a 2-photon microscope (Olympus FV1000 Confocal Microscope with MaiTai Deep Sea Laser) for autofluorescent imaging using a 4x objective (UPlanFL N 4x/0.13). NAD(P)H and FAD were excited at 800 nm (8, 9), to excite both simultaneously, and collected using a 505 nm dichroic mirror and a 460-500 nm filter, for NAD(P)H, or a 520-550 nm filter, for FAD. Separately, GFP was excited at 488 nm and collected. Z-stack images up to 1 mm deep at a 10 µm interval were obtained for image analysis. To analyze the images, we used a custom script in ImageJ (10).

Primary Cell Culture

Mouse lung fibroblasts were isolated using FACS as detailed above and plated in 6-well dishes (100,000 cells per well) with DMEM + 10% FBS + 1% anti-anti. After 5 days, the media was replaced for the first time and then every 2-3 days following. At the indicated timepoints, cells were lifted using 0.25% trypsin and suspended in autoMACS for FACS isolation and flow cytometry analysis.

qPCR

The mRNA of FACS-isolated cells was isolated using a RNeasy Micro Kit (Qiagen) according to manufacturer's directions and then converted into cDNA using Superscript Vilo cDNA Synthesis Kit (Invitrogen). qPCR of cDNA was performed using FastStart Essential DNA Green Master (Roche) and primers (Supp Table 2).

Fibrosis evaluation

Sections (5 µm thick) were cut from formalin-fixed paraffin-embedded lung tissues, and the sections were stained either with H&E or with Masson's Trichrome Stain Kit. Images were taken with an Olympus CKX53 or a Motic EasyScan for H&E or MTC, respectively.

Hydroxyproline content was measured using a hydroxyproline assay kit. Briefly, right lower lung and accessory lobe samples were transferred into glass tubes and hydrolyzed with 200 µl 6N HCL at 110°C for 48 h. The hydrolyzed samples were evaporated to remove excess HCL, reconstituted with 400 µl $H_2O$ and filtered in 1.5 ml centrifuge tubes equipped with a 0.45 µm semipermeable membrane filter. After samples were added to a 96 well micro-plate, Chloramine T solution was added, and the plate was incubated at room temperature for 20 min. 100 µl of Erlich's reagent was added to each well and the plate was incubated at 65°C for 18 min. OD 550 nm was obtained and compared to a hydroxyproline standard curve.

<u>Statistics</u>
Statistics were not performed on intravital imaging results due to low sample numbers. All other data are presented as the mean ± SEM. Statistical analysis for each figure is provided in each figure caption.

| **Supplemental Table 1.** Materials used in these studies | | | |
|---|---|---|---|
| **Product** | **RRID** | **Vendor** | **Produce Number** |
| Bleomycin | | Vizient Inc | 0409-0332-20 |
| 20 ga catheter, Angiocath | | BD | 26742 |
| Microsprayer | | PennCentury | IA-1C and FMJ-250 |
| PBS | | Life Technologies | 14190250 |
| Liberase-TM | | Sigma-Aldrich | 5401127001 |
| DNase | | Sigma-Aldrich | 4536282001 |
| 40 µm filter | | Sigma | SCNY00040 |
| RBC Lysis buffer | | BioLegend | 420301 |
| 30 µm filter | | SYSMEX AMERICA INC | NC9682496 |
| AutoMACS Buffer | | Miltenyi Biotec | 130-091-222 |
| CD45 MicroBeads, mouse | RRID:AB_2877061 | Miltenyi Biotec | 130-052-301 |
| LS Columns | | Miltenyi Biotec | 130-042-401 |
| Brilliant Violet 785(TM) anti-mouse CD31 | RRID:AB_2810334 | Biolegend | 102435 |
| Brilliant Violet 711(TM) anti-mouse CD45 | RRID:AB_2564383 | Biolegend | 103147 |
| Brilliant Violet 421(TM) anti-mouse CD326 (Ep-CAM) | RRID:AB_2563983 | Biolegend | 118225 |
| CD45 MicroBeads, human | RRID:AB_2783001 | Miltenyi Biotec | 130-045-801 |
| Brilliant Violet 421(TM) anti-human CD326 (Ep-CAM) | RRID:AB_2563847 | Biolegend | 324220 |
| Brilliant Violet 711(TM) anti-human CD45 | RRID:AB_2563466 | Biolegend | 304050 |
| Brilliant Violet 785(TM) anti-human CD31 | RRID:AB_2860782 | Biolegend | 303148 |
| TO-PRO™-3 Iodide | | Thermo Fisher Scientific | T3605 |
| Magic Red Cathepsin K Assay Kit | | Immunochemistry | 940 |
| Depilatory cream | | Nair | 22600223191 |
| Suture | | ThermoFisher | 50-118-0848 |
| Cyanoacrylate | | Fisher Scientific | NC0632797 |
| Coverslip | | Electron Microscopy Sciences | 72296-05 |
| 6-well tissue culture plates | | Corning | 353046 |
| DMEM | | Life Technologies | 11965118 |
| FBS | | ATCC | 30-2020 |
| Anti-anti | | Fisher Scientific | 15-240-062 |
| Trypsin-EDTA | | Sigma Aldrich | T4049-100ML |
| Rneasy Micro Kit | | Qiagen | 74004 |
| Superscript Vilo cDNA synthesis kit | | Invitrogen | 11754050 |
| FastStart Essential DNA Green Master | | Roche | 6924204001 |
| Hydroxyproline | | Biovision | K226 |
| Masson's Trichrome | | Polysciences, Inc. | 25088-1 |
| **Equipment** | | **Manufacturer** | **Model** |
| FACS Aria II | | BD | Aria II |
| LSR Flow Cytometer | | BD | Fortessa X-20 |
| Confocal Microscope w/ MaiTai DeepSea Laser | | Olympus | FV1000 |

| | | | |
|---|---|---|---|
| Confocal Objective | | Olympus | UPlanFL n 4x/0.13 |
| PCR machine | | Roche | Lightcycler 96 |
| Small animal ventilator | | Harvard Apparatus | 845 |
| **Software** | **RRID** | **Distributor** | **Software (version)** |
| Flow cytometry analysis software | RRID:SCR_008520 | BD | FlowJo (v10.8) |
| Statistical analysis software | RRID:SCR_002798 | GraphPad | Prism (v9.4) |
| Image analysis software | RRID:SCR_002285 | ImageJ | Fiji (v1.53t) |

Supplemental Table 2. Primers used in these studies.

| | Forward | Reverse | Reference |
|---|---|---|---|
| **Gusb** | GGCTGGTGACCTACTGGATTT | GGCACTGGGAACCTGAAGT | (11) |
| **Nox4** | CGGGATTTGCTACTGCCTCCAT | GTGACTCCTCAAATGGGCTTCC | (12, 13) |
| **Ptgs2** | GCGACATACTCAAGCAGGAGCA | AGTGGTAACCGCTCAGGTGTTG | (14, 15) |
| **Col1a1** | ATCATAGCCATAGGACATCTGG | CTGGACAGCCTGGACTTC | (16, 17) |
| **Fn1** | CCAGCAGCATGATCAAAACAC | GGTGGCTACATGTTAGAGTGTC | (17) |
| **Mmp14** | GGATGGACACAGAGAACTTCG | TTTTGGGCTTATCTGGGACAG | (18, 19) |
